# Purification, crystallization, and preliminary structural analysis of multivalent immunogenic effector protein-anchored SARS-CoV-2 RBD

**DOI:** 10.1101/2023.08.17.553661

**Authors:** Taek Hun Kwon, Tae Gyun Kim

## Abstract

The continuous spread of highly transmissible variants of concern and the potential diminished effectiveness of existing vaccines necessitate ongoing research and development of new vaccines. Immunogenic molecule-anchored antigen has demonstrated superior efficacy in subunit vaccination, primarily due to enhanced cellular uptake facilitated by the affinity between the surface of Immunogenic molecule and the cell membrane. Based on the Immunogenic recombinase *B. malayi* RecA (*Bm*RecA), we have overexpressed the construct of *Bm*RecA with SARS-CoV-2 RBD (*Bm*RecA-RBD) that exists as a stable helical filament formation; it was purified and crystallized to obtain X-ray diffraction data at 2.7 Å, belonged to the hexagonal symmetry group *P*6_5_ in the unit-cell parameters of a=b=122.12, c=75.55 and α=β=90°, γ=120°. The Matthews coefficient was estimated to be 3.12 Å^3^ Da^-1^, corresponding to solvent contents of 52.65.

## INTRODUCTION

Highly transmissible respiratory diseases such as COVID-19 present significant threats to global public health. The continuous spread of extremely transmissible variants of concern like alpha, beta, gamma, delta, and omicron has prompted apprehension regarding the efficacy of existing COVID-19 vaccines, specially in terms of preventing breakthrough infections (Ahmad *et al*., 2022). Furthermore, the effectiveness of currently available vaccines in protecting vulnerable populations, particularly those who are immunocompromised or elderly, may be diminished. This emphasizes the importance of continuous research and the development of new vaccines to effectively address this concern (Thye *et al*., 2021). The immunogenic molecule-anchored antigens outperform monomeric, soluble antigens in terms of efficacy when it comes to subunit vaccination (Smith *et al*., 2013). The enhanced cellular uptake of anchored antigen, in comparison to antigen alone, can be attribute to the increased affinity between the surface of immunogenic effector protein and the cell membrane (Lung *et al*., 2020; Rodrigues *et al*., 2021). Several proteins have recently emerged as a promising platform for immunogenic antigens in condition of monodispersity, biocompatibility, biodegradability, low-cost productivity, and a hollow cavity capable of reversible assembly/disassembly (Khoshnejad *et al*., 2018).

Prokaryotic RecA and eukaryotic Rad51 structures have been reported as helical nucleoprotein filament on the oligonucleotide (ssDNA/dsDNA). Based on structural and functional comparisons of key enzymes in the repairing damaged DNA, resolving DNA replication errors and promoting genetic diversity. The recombinase protein of Wolbachia from *Brugia malayi* (*Bm*RecA) has the significant sequence which shows structural similarities among other RecAs and promotes immune-reactivities such as significant cellular adherence and cytotoxicity (Gangwar *et al*., 2019). The amino acid sequence of *Brugia malayi* (Uniprot # Q5GSK9) shows the average sequence identity of 51% and 12% to that of well-known RecA *Escherichia coli* (Uniprot # P0A7G6) and Rad51 from *Homo sapiens* (Uniprot # Q06609) (Fig. 2).

**Figure 1.**
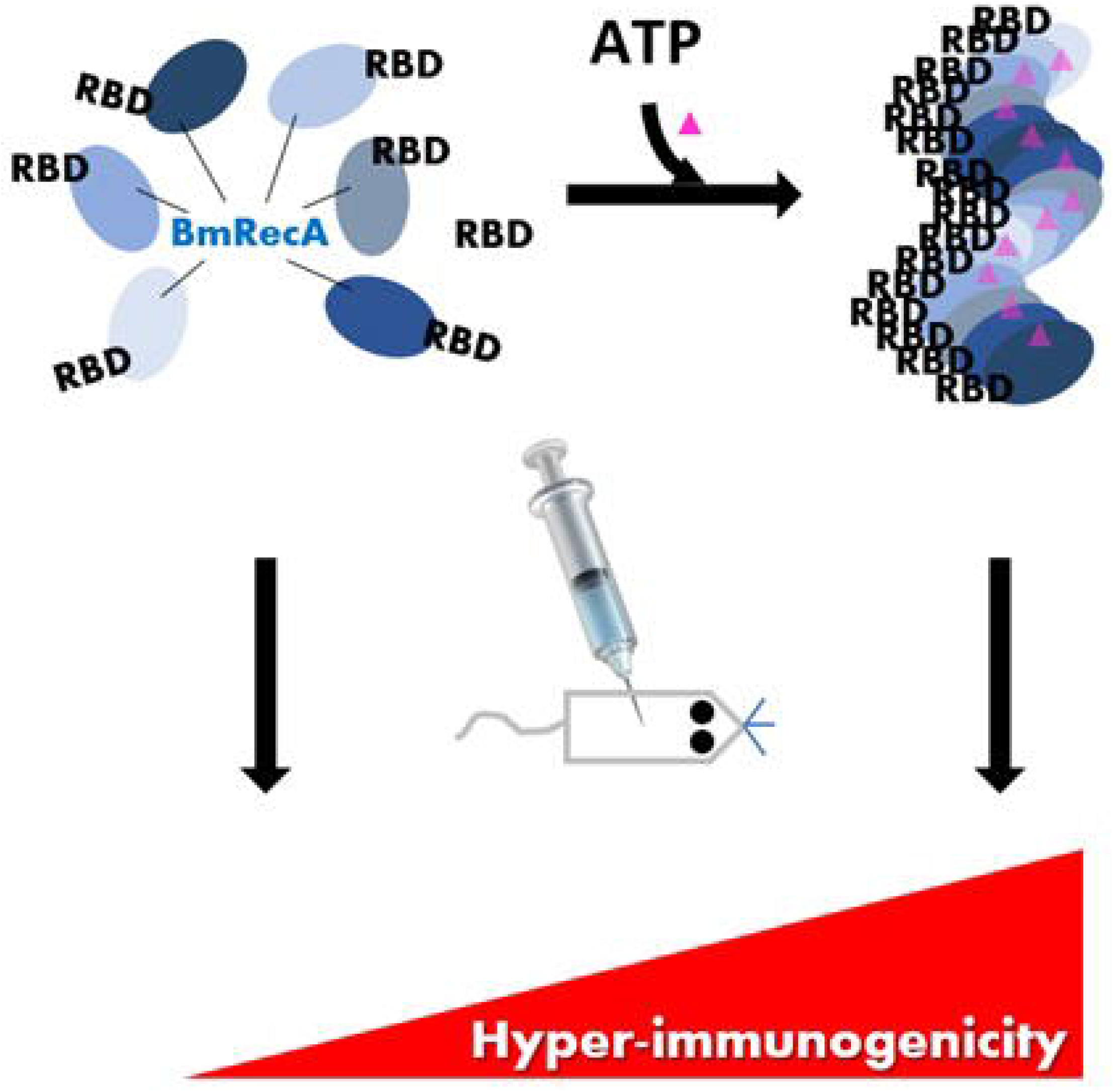
Proposal scheme on the antigen immunogenicity between monomer and multimeric *Bm*RecA-RBDs.

**Figure 2.**
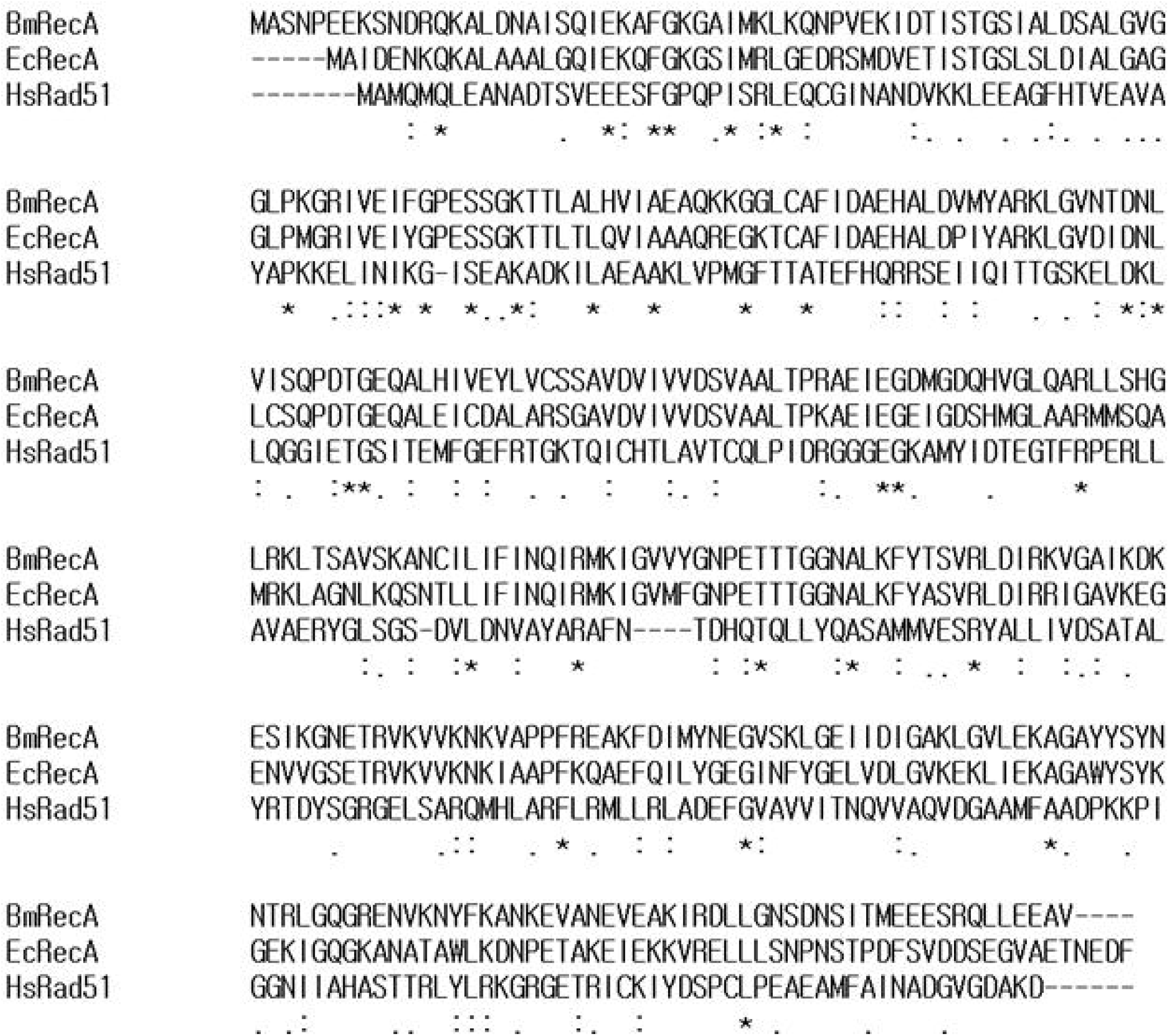
The amino acid sequence alignment among *B. malayi* RecA, *E. coli* RecA and *H. sapiens* Rad51 was generated and modified by multiple sequence alignment by ClustalW (www.genome.jp/tools-bin/clustalw). The accession number and database of aligned sequences are indicated in parentheses: BmRecA, *Brugia malayi* (Uniprot # Q5GSK9); EcRecA, *Escherichia coli* (Uniprot # P0A7G6); HsRad51, *Homo sapiens* (Uniprot # Q06609).

To date, immunogenic molecule-mediated antigens have been identified as assemblies of polypeptides that present multivalent subunit-based antigen. Here, we prepare the construct of *Bm*RecA with SARS-CoV-2 RBD epitope (*Bm*RecA-RBD) and preliminarily analysed the structural solution on *Bm*RecA-RBD at high resolution using X-ray crystallographic and negative-stain electron microscopic techniques. Reproducible crystals for *Bm*RecA-RBD were obtained and diffracted to 2.7 Å resolution. Thus, this study will show structural features and molecular characteristics of RBD construct with highly immunogenic *Bm*RecA sequence as a putative vaccine candidate.

## RESULTS AND DISCUSSION

The codon-optimized *Bm*RecA-RBD-encoding DNA was synthesized using commercially chemical synthesis (IDT, Singapore). The *Bm*RecA-RBD was cleaved by using *BamH*I and *Xho*I restriction enzymes and subcloned into the pET-28a-TEV expression plasmid (Table 1). When the protein was initially transformed into *E. coli* BL21(DE3), expression level of *Bm*RecA-RBD was low. After changing culture temperature and adding 0.5% glucose in LB culture media, the *Bm*RecA-RBD expression showed a significant increase to be obtained as a soluble fraction. The recombinant *Bm*RecA-RBD protein with an N-terminal TEV protease-cleavable His_6_-tag was successfully overexpressed as a soluble form in *E. coli* BL21(DE3) and purified to crystallizable quality, followed Ni-NTA affinity chromatography, cleavage of the N-terminal His-tag with TEV protease, and Superose 6 increase 10/300 GL (Cytiva, USA). The final yield of *Bm*RecA-RBD protein was concentrated at approximately 2 mg per L of cultured cells, and the highly qualified purity of the protein sample was judged by Coomassie-stained SDS-PAGE, indicating as a monomeric 64.1 kDa calculated using the ExPASy server (Wilkins *et al*., 1999) (Fig. 3A). Fractions containing *Bm*RecA-RBD were collected and concentrated to 12 mg/ml for crystallization.

**Table 1.**
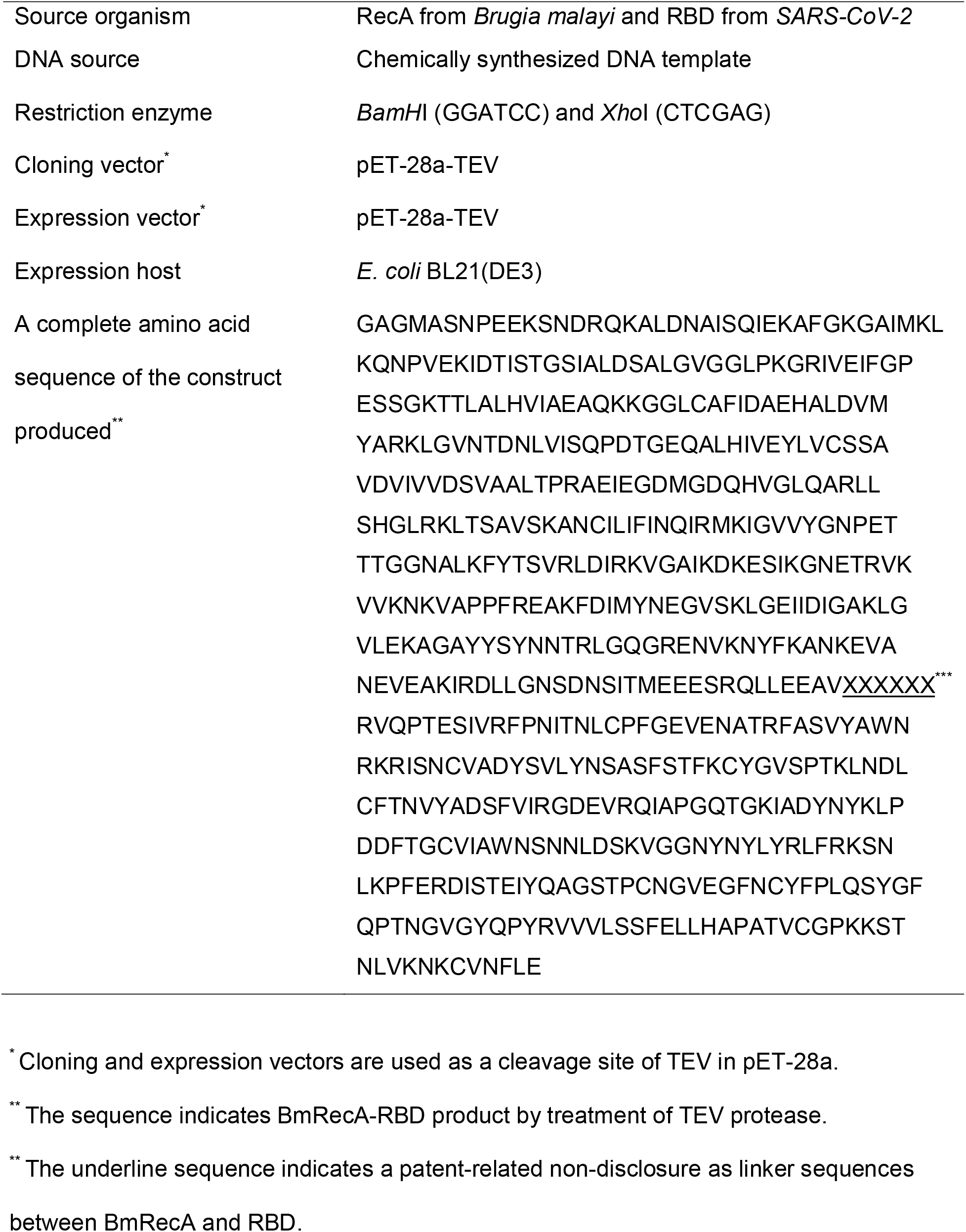
Macromolecule production information.

**Figure 3.**
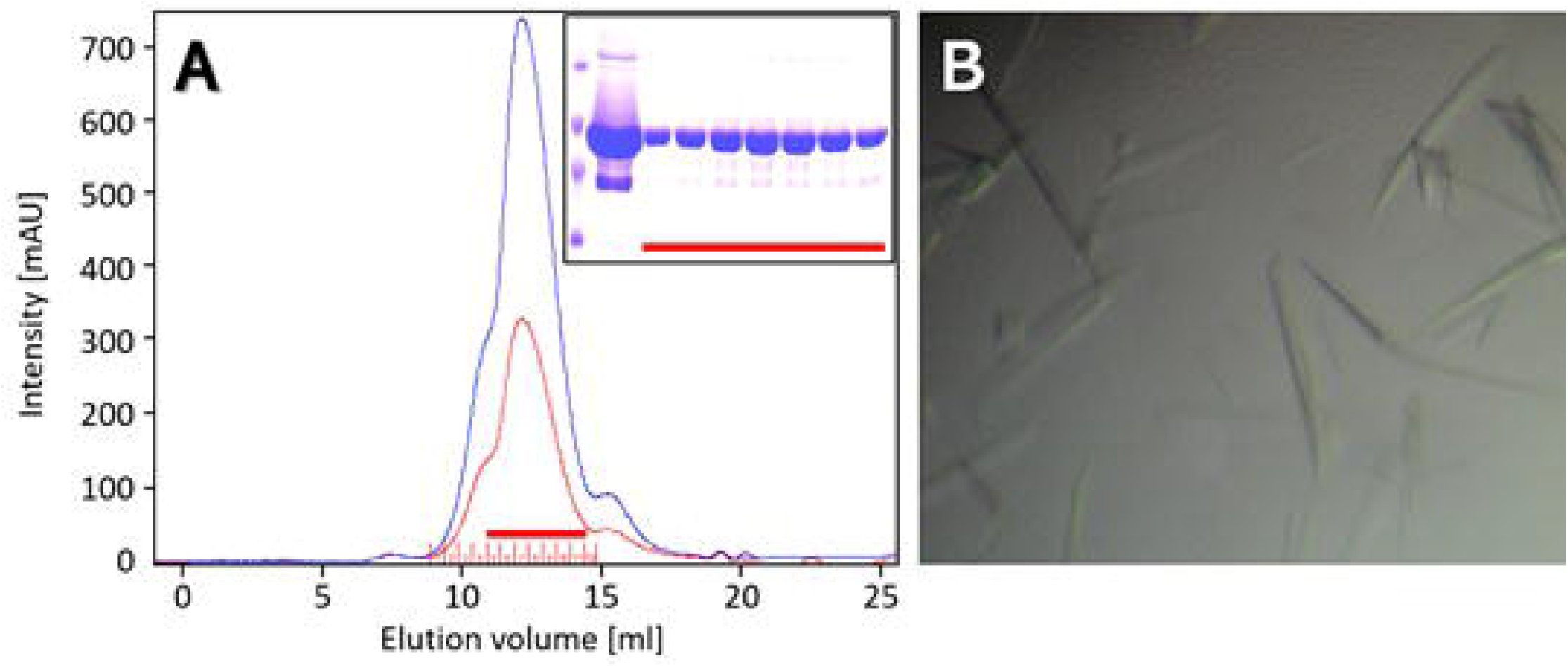
Purification and crystallization of *Bm*RecA-RBD. (A) Size exclusion chromatography profile and SDS-PAGE analysis of *Bm*RecA-RBD. Inlet SDS-PAGE shows the molecular weight of standard size markers indicated as 97, 66, 45, and 30 kDa (up-to-down). The underline was collected and then concentrated to 12 mg/ml for crystallization. (B) The *Bm*RecA-RBD crystals were obtained as blade-shape forms with the following condition of 0.05 *M* Hepes-KOH, pH 7.3, 20% (w/v) MPD, 0.1 *M* Mg(OAc)_2_.

Initial crystallization of *Bm*RecA-RBD was carried out using the sitting-drop vapour diffusion method at 20°C. For diffraction-quality crystals, *Bm*RecA-RBD was obtained using 0.05 *M* Hepes-KOH, pH 7.3, 15% (w/v) MPD, 0.1 *M* MgCl_2_, indicating that the concentration of MPD was critical to be crystallized as an optimized formation *Bm*RecA-RBD. Within one week, the blade-shaped crystals of *Bm*RecA-RBD appeared in the average crystal size being 0.45 × 0.07 × 0.05 mm (Fig. 3B). The optimized crystals were obtained in the presence of 0.05 *M* Hepes-KOH, pH 7.3, 20% (w/v) MPD, 0.1 *M* Mg(OAc)_2_ (Table 2). After transferring *Bm*RecA-RBD crystals into additional 30% (v/v) glycerol in mother liquor for cryo-protection, the optimized *Bm*RecA-RBD crystals were directly flash-cooled in a gas stream from liquid N_2_. Crystals belonged to the hexagonal symmetry space group *P*6_5_ with three monomers per asymmetric unit. The Matthews coefficient for *Bm*RecA-RBD was estimated to be 3.12 Å^3^ Da^-1^, corresponding to solvent contents of 52.65% (Matthews, 1968).

**Table 2.**
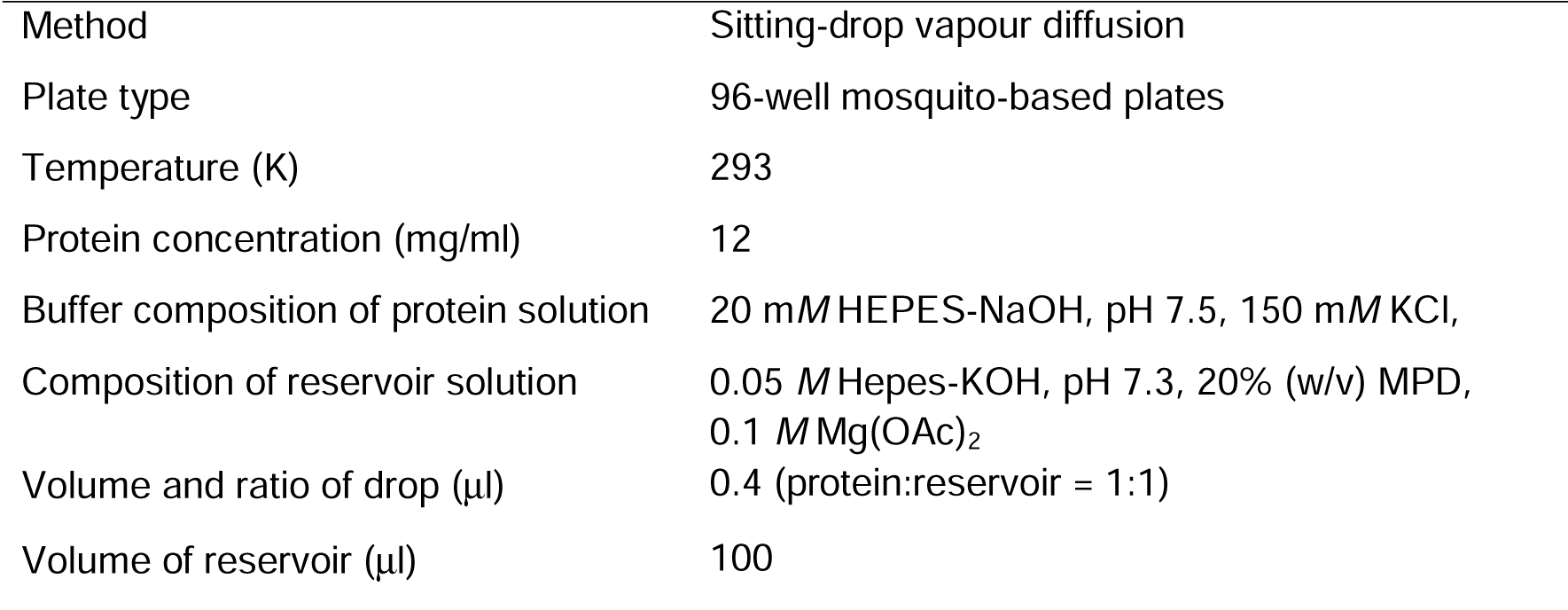
Protein crystallization

|  |  |
| --- | --- |
| Method | Sitting-drop vapour diffusion |
| Plate type | 96-well mosquito-based plates |
| Temperature (K) | 293 |
| Protein concentration (mg/ml) | 12 |
| Buffer composition of protein solution | 20 mM HEPES-NaOH, pH 7.5, 150 mM KCl, |
| Composition of reservoir solution | 0.05 M Hepes-KOH, pH 7.3, 20% (w/v) MPD,<br>0.1 M Mg(OAc)<sub>2</sub> |
| Volume and ratio of drop (μl) | 0.4 (protein:reservoir = 1:1) |
| Volume of reservoir (μl) | 100 |

The molecular replacement was performed with CCP4 MolRep (Winn et al., 2011) and PHASER (McCoy et al., 2007) using the *E. coli* RecA structure (PDB code 4TWZ) as a search model. Initial refinement produced a possible model of *Bm*RecA-RBD with R_work_ of 28.4% and R_free_ of 34.1% using PHENIX (Adams *et al*., 2010). To interpret the electron density and the phase information, molecular replacement was carried out done using CCP4 MolRep (Winn et al., 2011) and PHASER (McCoy et al., 2007). Preliminary structure refinement and manual build-up of the atomic models were performed using PHENIX:refine (Adams *et al*., 2010) and WinCOOT program (Emsley and Cowtan, 2004). Further model building and refinement of *Bm*RecA-RBD structure was of well-qualified map to build a three-dimensional structure for *Bm*RecA-RBD crystals diffracted to a resolution of 2.7 Å. Statistics for diffraction data collection and processing details are summarized in Table 3.

**Table 3.**
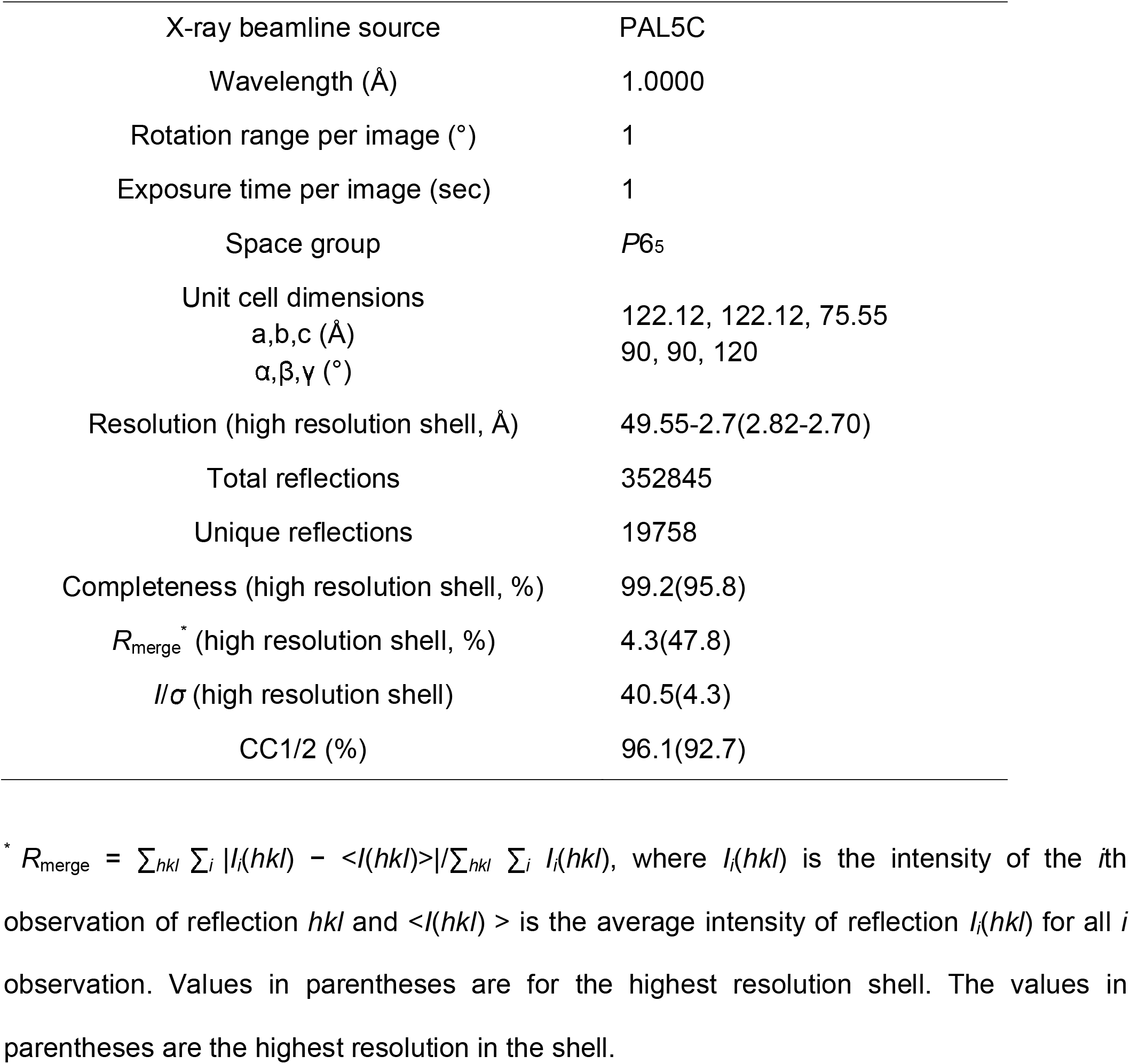
Data collection and processing statistics

To investigate a molecular distribution pattern for *Bm*RecA-RBD protein using negative-stain transmission electron microscopic (TEM) method, we showed the structural morphology of nucleoside- and ssDNA-bound *Bm*RecA-RBD proteins (Fig. 4A & 4B, respectively), indicating that conserved known *Bm*RecA-RBD structures has a helical filamentous formation like bacterial RecA (Hertzog *et al*., 2023).

**Figure 4.**
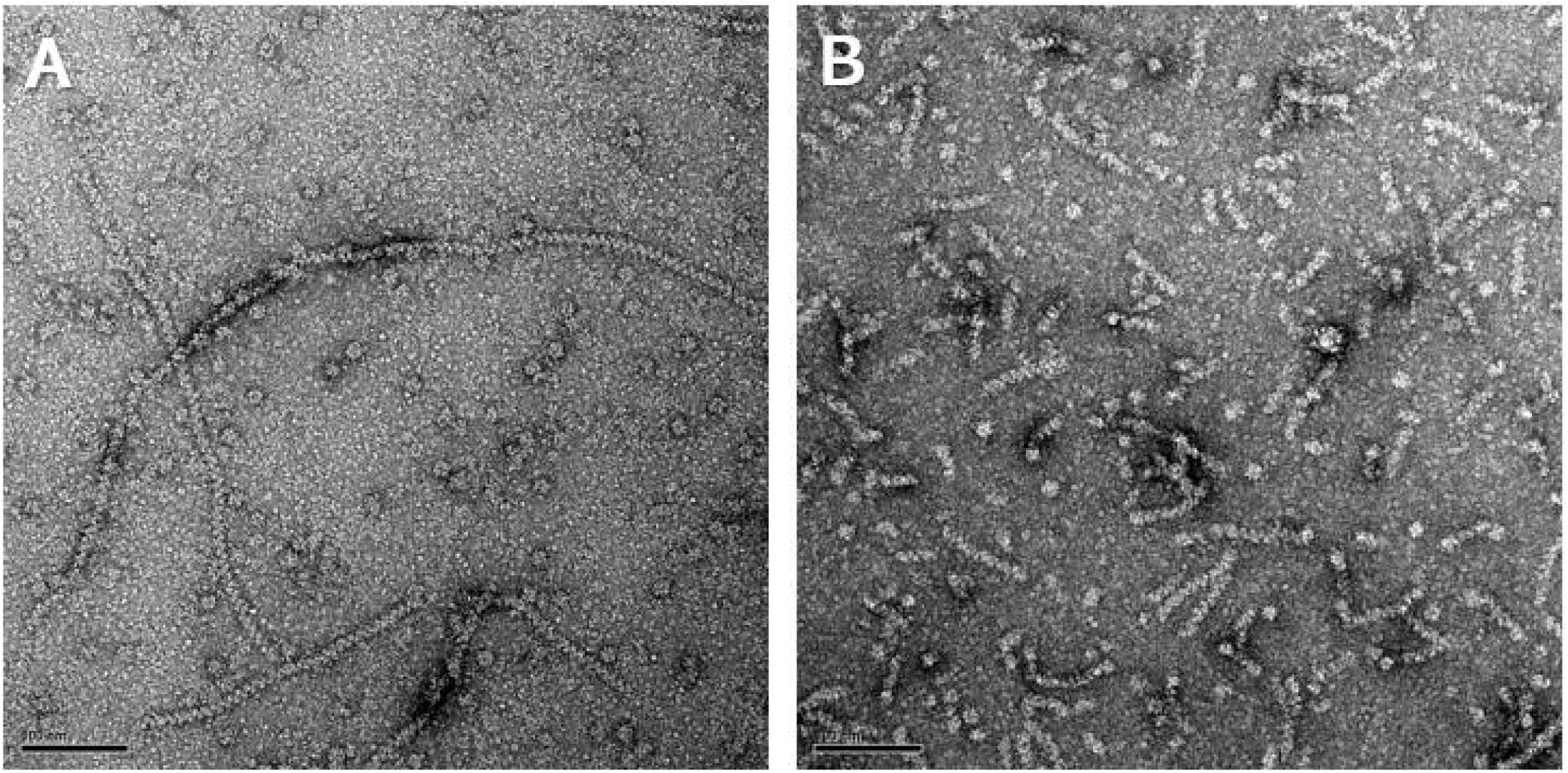
Negative-stain TEM analysis of ATPgS-bound *Bm*RecA-RBD (A) and ATPgS-bound *Bm*RecA-RBD (B) in the presence of 30mer ssDNA. The scale bar indicates 100 nm.

In this study, the purification, crystallization, X-ray crystallography and negative-stain TEM analysis provide a structural investigation of *Bm*RecA-RBD to explain the further determination of flexible structural features compared to the current *Bm*RecA-RBD structure and the mechanical understanding antigen RBD construct with highly immunogenic *Bm*RecA sequence as a putative vaccine candidate.

## METHODS

### Protein expression and purification

Commercial synthesis of *Bm*RecA-RBD encoding was used as a chimeric construct shown in Table 1. To confirm restriction enzyme sites that are excluded in the full coding sequence of *Bm*RecA-RBD, the uncleaved restriction enzyme site was checked with NEBcutter V2.0. We selected *Bam*HI and *Xho*I restriction sites of the expression vector pET-28a-TEV (Tobacco Etch Virus), which includes a cleavage site by TEV protease site in the N-terminus. To carry out to over-expressed cells, the pET-28a-TEV-*Bm*RecA-RBD was transformed and expressed in *E. coli* BL21(DE3) (Invitrogen, USA) cells within 0.5% glucose-mediated LB media at 23°C, and induced with 0.4 m*M* Isopropyl-D-thiogalactoside (Sigma, USA) to over-express the recombinant proteins. After collection of cultured cells by centrifugation, cell pellets were resuspended with buffer A (20m*M* Tris-HCl, pH 8.0, 300 m*M* NaCl, 5m*M* β-mercaptoethanol, 1mg/ml lysozyme, and mixture of protease inhibitors included in aprotinin, leupeptin, pepstatin, and PMSF) and gently disrupted by VC505 sonicator (Sonics, USA) on ice. Cell debris and supernatant were separated by high-speed centrifugation at 20,000 x g for 1 h at 4 °C. Additional impurities from collected supernatant were filtrated by using 0.45 μm MF-Millipore Membrane filter (Merck, USA). The filtered supernatant included *Bm*RecA-RBD protein was applied to a nickel-nitrilotriacetic acid (Ni-NTA, Cytiva) affinity column pre-equilibrated with buffer A. The Ni-NTA column was washed by using 20 times of the column volume by buffer A and 50m*M* Imidazole. Target protein was eluted with a linear gradient using buffer B (50 m*M* Tris-HCl, pH 8.0, 300 m*M* NaCl, 300 m*M* imidazole, and 5 m*M* β-mercaptoethanol). Eluted *Bm*RecA-RBD protein was dialyzed against buffer A while simultaneously digesting with TEV protease for at least 12 hrs at 4 °C to cleave the His-tagging peptide. To obtain the cleaved *Bm*RecA-RBD protein through re-application into Ni-NTA column, the TEV-treated *Bm*RecA-RBD was concentrated using Amicon centrifugal filters (Millipore, USA) with a cut-off of 10 kDa and injected into Superose 6 increase 10/300 GL (Cytiva, USA) equilibrated with buffer C (20 m*M* Hepes-NaOH, pH 7.5, 150 m*M* KCl). *Bm*RecA-RBD protein fractions were examined by sodium dodecyl sulfate-polyacrylamide gel electrophoresis (SDS-PAGE) and concentrated at 12 mg/ml to use crystallization (Fig. 3A). Protein preparation is shown in Table 1.

### Crystallization

*Bm*RecA-RBD protein was initially crystallized using various screening kits from commercial company as Hampton Research, Creative Biostructure, and Jena Bioscience using an automatic crystallization machine Mosquito LCP (SPT Labtech, USA). Based on the initial screening, both were further optimized as diffraction-qualified crystals using 96-well plates handled with the sitting-drop vapour diffusion method as same condition in Mosquito LCP machine applied with varying concentrations of buffer, precipitant, and metal ion. *Bm*RecA-RBD crystals were grown at 293 K in 1 μL hanging drops comprised of equal volumes of protein stock solution (12 mg/mL) and mother liquor containing 0.05 *M* Hepes-KOH, pH 7.3, 20% (w/v) MPD, 0.1 *M* Mg(OAc)_2_ (Fig. 3B). Crystals grew to a maximum size of 0.45 × 0.07 × 0.05 mm in one week. Crystals were transferred into cryoprotectant (30% (*v*/*v*) glycerol in reservoir solution) and were frozen in liquid N_2_ at near 77K. Crystallization information is summarized in Table 2.

### Data collection and initial processing

*Bm*RecA-RBD crystals were harvested using cryo-loops and immersed promptly in cryo-protectant solution. X-ray diffraction data sets for apo and *Bm*RecA-RBD crystals were collected at 2.7 Å resolution on beamline 5C at the Pohang Accelerator Laboratory, Republic of Korea. Exposure time was 1.0 s per frame. One complete data set was obtained with 1° of oscillation angle. X-ray diffraction data were collected at 100K. The diffraction datasets were indexed, merged and scaled with *HKL*2000 (Otwinowski *et al*., 1997). Crystallographic data statistics are presented in Table 3. The molecular replacement search was performed by using CCP4 MolRep (Winn et al., 2011) and PHASER (McCoy et al., 2007). Data collection and processing statistics are presented in Table 3.

### Negative-stain TEM analysis

For preparing *Bm*RecA-RBD on carbon EM grid (EMJapan, Japan), we prepared one carbon EM grid and place on the glass slide with the carbon side up (dark side of the grid up) using the fine tweezers and then placed the glass slide containing the EM grid in the PELCO easiGlow (Ted Pella, USA). Treated carbon EM grid by the self-closing tweezers were used to protein solution. The 1 mM ATPgS- and 30 μM ssDNA-bound *Bm*RecA-RBD proteins were diluted to a final concentration ranging from 0.015 to 0.05 mg/ml were placed onto freshly glow-discharged continuous carbon-coated side of carbon EM grids, stained with final 1% uranyl acetate for 30 sec, blotted by gently removing the edge of grid to filter paper, and then completely air-dried for 1h. Micrographs were recorded in a JEM-1230R transmission electron microscope operated at 100 kV using Gatan Ultracan digital camera at a nominal magnification of 30,000.

## ACKNOWLEDGEMENTS

We appreciate the technical assistance of beamline 5C at Pohang Accelerator Laboratory for help with X-ray diffraction and data collection experiments. We gratefully acknowledge direct financial support from Gyeongbuk Institute for Bio industry and the Ministry of Trade, Industry, and Energy (P0009787).

